# An RNA modification enzyme directly senses reactive oxygen species for translational regulation in *Enterococcus faecalis*

**DOI:** 10.1101/2022.10.12.511899

**Authors:** Wei Lin Lee, Ameya Sinha, Ling Ning Lam, Hooi Linn Loo, Jiaqi Liang, Peiying Ho, Liang Cui, Cheryl Siew Choo Chan, Thomas Begley, Kimberly Kline, Peter Dedon

## Abstract

Bacteria possess elaborate systems to manage reactive oxygen and nitrogen species (ROS) arising from exposure to the mammalian immune system and environmental stresses. Here we report the discovery of an ROS-sensing RNA-modifying enzyme that regulates translation of stress-response proteins in the gut commensal and opportunistic pathogen *Enterococcus faecalis*. We analyzed the tRNA epitranscriptome of *E. faecalis* in response to reactive oxygen species (ROS) or sublethal doses of ROS-inducing antibiotics and identified large decreases in N^2^-methyladenosine (m^2^A) in both 23S ribosomal RNA and transfer RNA. This we determined to be due to ROS-mediated inactivation of the Fe-S cluster-containing methyltransferase, RlmN. Genetic knockout of RlmN gave rise to a proteome that mimicked the oxidative stress response, with increased levels of superoxide dismutase and decreased virulence proteins. While tRNA modifications are established to be dynamic for fine-tuning translation, here we report the first instance of a dynamically regulated, environmentally responsive rRNA modification. These studies lead to model in which RlmN serves as a redox-sensitive molecular switch, directly relaying oxidative stress to modulating translation through the rRNA and the tRNA epitranscriptome, revealing a new paradigm for understanding direct regulation of the proteome by RNA modifications.

## Main Text

Bacteria possess elaborate systems to manage reactive oxygen species (ROS) arising from exposure to the mammalian immune system and environmental stresses. Here we report the discovery of an ROS-sensing RNA-modifying enzyme that regulates translation of stress-response proteins in the gut commensal and opportunistic pathogen *Enterococcus faecalis*. Following exposure to the superoxide generator, menadione, or sublethal doses of ROS-inducing erythromycin and chloramphenicol, analysis of 24 modified ribonucleosides of the epitranscriptome revealed large decreases in N^2^-methyladenosine (m^2^A) in both 23S ribosomal RNA and transfer RNA caused by ROS-mediated inactivation of the Fe-S cluster-containing methyltransferase, RlmN. Loss of RlmN altered protein expression in a way that mimicked menadione exposure, such as increased superoxide dismutase and decreased virulence proteins. These studies suggest that RlmN acts as a redox-sensitive molecular switch that links environmental and antibiotic-induced ROS exposure to epitranscriptome dynamics in ribosomal and transfer RNA to effect translation of stress response proteins.

While transcriptional regulation in response to stress is well established in bacteria, translational regulation is less well understood. We recently demonstrated that hypoxic stress in mycobacteria caused reprogramming of dozens of RNA modifications in the tRNA epitranscriptome, which was linked to codon-biased translation of stress response transcripts^1^. Hypothesizing that the same would be true for *Enterococcus faecalis*, we asked how the stress of antibiotic exposure would alter the levels of 24 modifications in the rRNA and tRNA. Here we examined two *E. faecalis* strains: OG1RF, a strain derived from the human commensal oral isolate OG1^5^, and V583^2^, a multidrug resistant clinical isolate. V583 possesses an erythromycin resistance methyltransferase (ErmB) that methylates the N^6^-position of adenosine (m^6^A, m^6,6^A) in 23S rRNA at position 2058 (*Escherichia coli* numbering)^3^ and prevents binding of macrolides (*e.g*., erythromycin), lincosamides, and streptogramin B (MLS)^3^, but only confers partial resistance to erythromycin^4^. OG1RF lacks ErmB and is thus about 100-fold more sensitive to erythromycin than V583.

We first explored antibiotic effects on V583’s epitranscriptome by growing cells in the presence of sub-inhibitory concentrations of erythromycin (10-200 **μ**g/mL; **Fig. 1A**). Following purification of 23S and 16S rRNAs and tRNA and hydrolysis to ribonucleosides, 24 modified ribonucleosides were quantified by chromatography-coupled mass spectrometry (LC-MS) in each type of RNA (**Fig. 1B-D, Supplementary Table 1**)^1^. Neither m^6^A nor m^6,6^A increased in 23S rRNA with erythromycin treatment, confirming that Erm expression in V583 is constitutive^4^. While most of the monitored modifications remained relatively unchanged with treatment, a striking reduction in 2-methyladenosine (m^2^A) in both 23S rRNA and tRNA was observed for erythromycin exposure (**Fig. 1B-D**), with dose-dependency (**Fig. 1E, F**).

**Figure 1.**
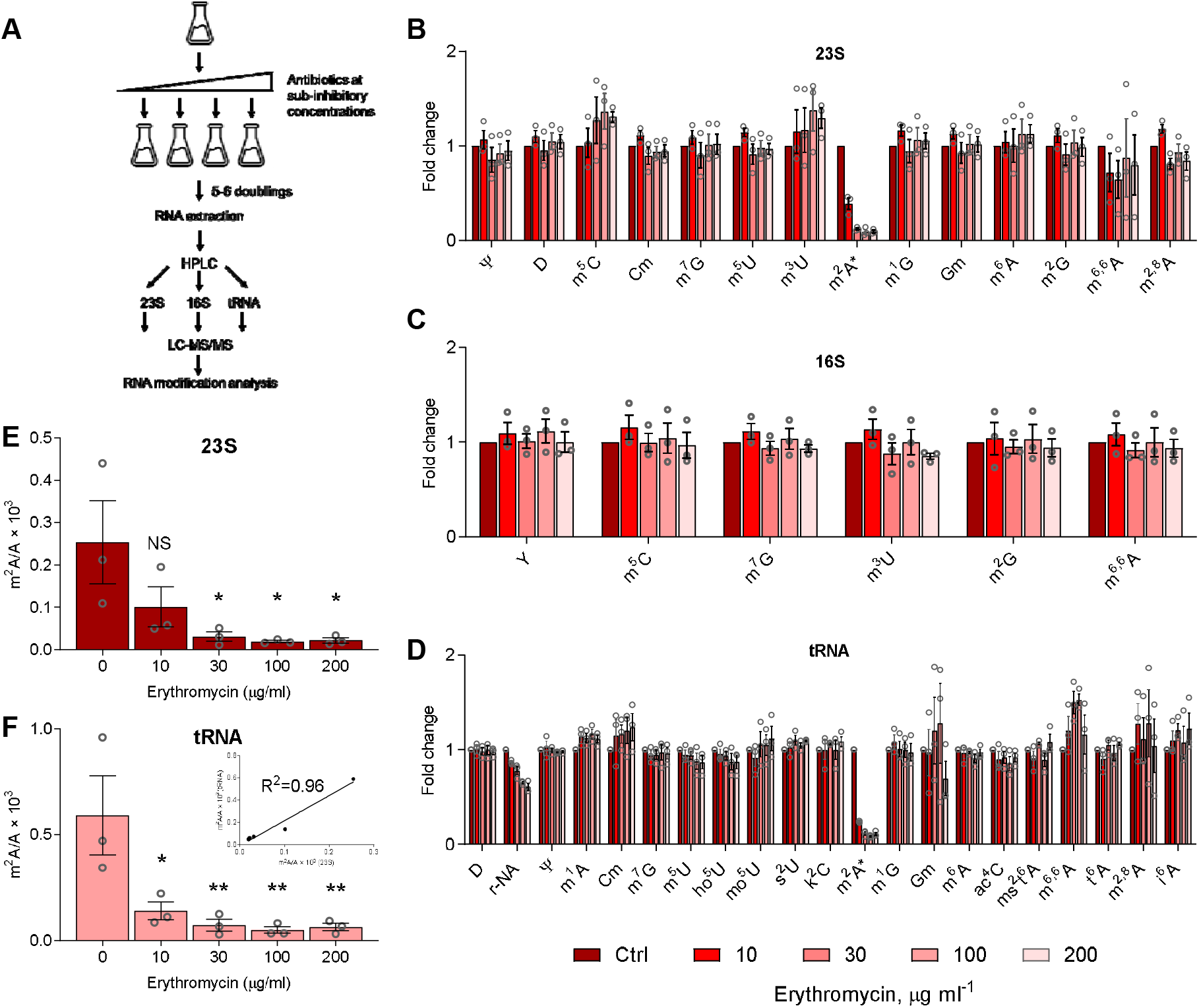
Epitranscriptomic profiling of 23S and 16S rRNA and tRNA of *Enterococcus faecalis* V583 grown in the presence of growth-permissive concentrations of erythromycin. (**A**) Epitranscriptome profiling workflow. Log-phase cultures of V583 were diluted with growth medium containing erythromycin below its MIC (10-200 μg/mL) and allowed to grow for 5-6 doublings to mid-log phase, after which RNA was isolated and RNA modifications quantified by LC-MS. (**B-D**) Changes in the levels of RNA modifications in V583 at varying doses of erythromycin (key at bottom of panel **D**) compared to untreated cells. Modification levels are shown as fold-change relative to an untreated control for 23S rRNA (**B**), 16S rRNA (**C**), and tRNA (**D**). Modifications are arranged from left to right in ascending retention time. *, m^2^A. See **Supplementary Table 1** for names, retention times, and precursor and product ion masses for the RNA modifications. (**E, F**) Effect of erythromycin dose on the ratio of m^2^A to adenosine. Ribonucleosides were quantified using LC-MS calibration curves, as illustrated for tRNA in the inset. All data are derived from 3 independent experiments (mean□±□SEM, *n*□=□3). Statistical analysis by one-way analysis of variance (ANOVA) with Dunnett’s test versus untreated controls: P<0.05 and P<0.005 are denoted as * and ** respectively.

To assess if the m^2^A reduction was unique to V583, we repeated the study with *E. faecalis* OG1RF at growth-permissive erythromycin concentrations (0.1-0.3 **μ**g/mL; **Supplementary Table 2**). We again observed a significant concentration-dependent decrease in m^2^A in both 23S rRNA and tRNA (**Fig. 2A, B**), with insignificant changes in the other 23 modifications (**Supplementary Fig. 2**). We next asked if m^2^A reduction was unique to erythromycin or a general response to all antibiotics. At 10-25% of the MICs of the different antibiotics (**Supplementary Table 2**), m A was reduced by macrolides erythromycin and spiramycin and the phenicol antibiotic chloramphenicol, all of which are bacteriostatic, but not by the bactericidal ciprofloxacin, ampicillin, or aminoglycosides kanamycin and gentamicin (**Fig. 2A-C**). The only mechanistic commonality here involves macrolide and chloramphenicol binding to the large (50S) ribosomal subunit at the peptide exit tunnel and peptidyl transferase center, respectively, while aminoglycosides target the small (30S) ribosomal subunit. This ribosome commonality stands in contrast to ciprofloxacin and ampicillin as gyrase inhibitor and cell wall disruptor, respectively^6^. The antibiotic-induced reduction of m^2^A is less in OG1RF compared to V583 (**Fig. 2C**), most likely due to the markedly lower concentrations of erythromycin and chloramphenicol used with OG1RF. So far, the data reveal that exposure of *E. faecalis* to sub-inhibitory concentrations of antibiotics that target the 50S ribosomal subunit selectively reduce m^2^A in 23S rRNA and tRNA, which raises the question of the mechanistic basis for this epitranscriptome behavior.

**Figure 2.**
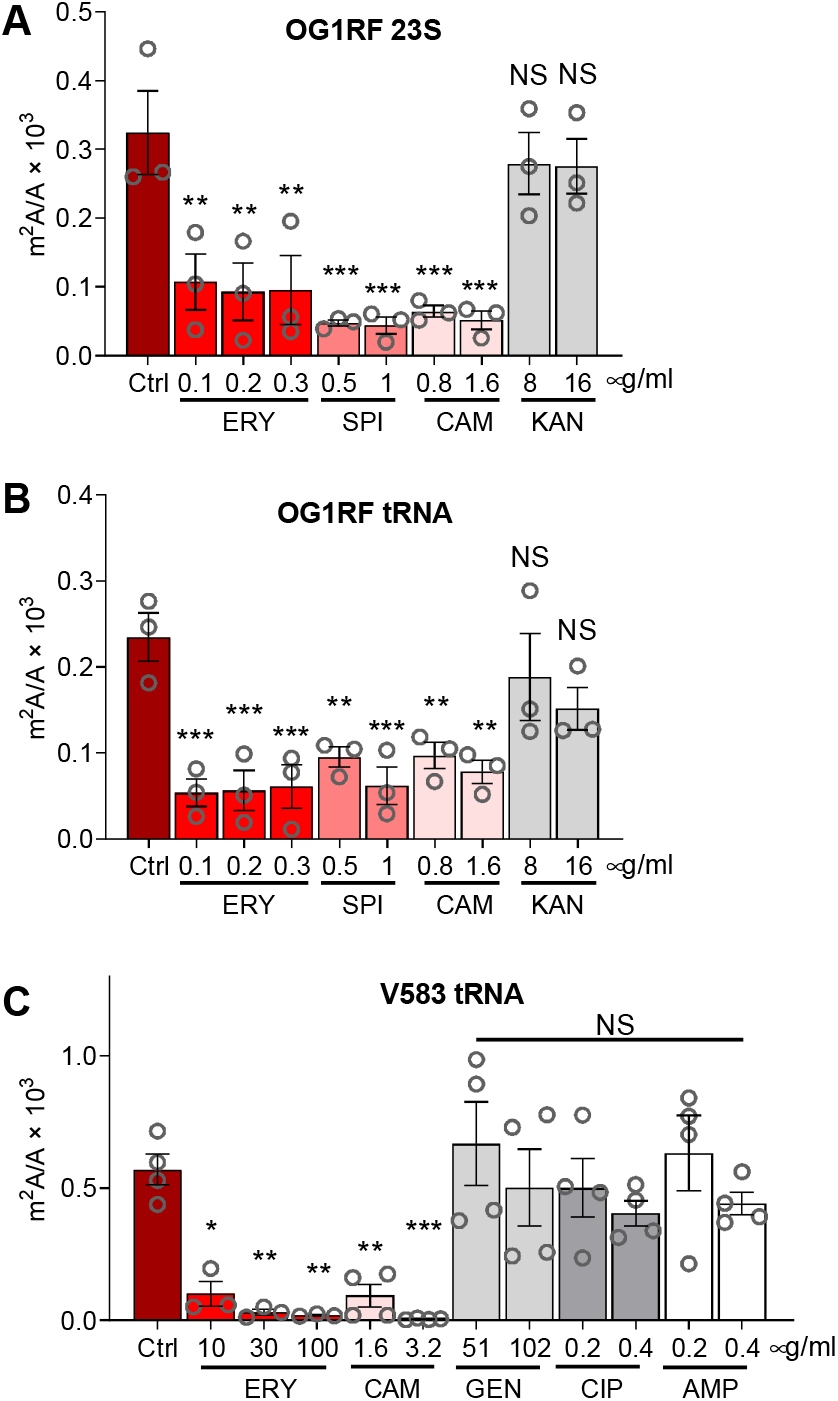
Reduction of m^2^A in V583 and OG1RF is specific to bacteriostatic macrolides and chloramphenicol. OG1RF (**A, B**) and V583 (**C**) were exposed to antibiotics at concentrations below their minimal inhibitory concentrations (MICs) and the quantity of m^2^A in tRNA and 23S rRNA measured by LC-MS. m^2^A levels decreased for the bacteriostatic macrolides erythromycin (ERY; **A, B, C**) and spiramycin (SPI; **B**) as well as chloramphenicol (CAM; **A, B, C**), but not for bactericidal ciprofloxacin (CIP; **C**), ampicillin (AMP; **C**), or the aminoglycosides gentamicin (GEN; **C**) and kanamycin (KAN; **A, B**). Data represent mean□±□SD, for n□=4 (**C**) and n=3 (**A, B**). Statistical analysis by one-way analysis of variance (ANOVA) with Dunnett’s test versus untreated controls. NS, not significant; P<0.05, P<0.005, and P<0.0005 are denoted as *, **, and ***, respectively.

In *Escherichia coli*, m^2^A formation is catalyzed by RNA methyltransferase RlmN, which methylates A2503 in 23S rRNA at the peptidyl transferase center in the 50S ribosomal subunit and A37 in the subset of tRNAs bearing adenine at this position in the anticodon stem loop^3^. Since RlmN in *E. faecalis* has not been previously characterized, we analyzed m^2^A levels in a *ΔrlmN* deletion mutant in OG1RF and found complete loss of the modification in 23S rRNA and tRNA (**Fig. 3A**). We first asked if the reduction of m^2^A by macrolides and chloramphenicol involved transcriptional or translational regulation of RlmN. Neither the level of *rlmN* mRNA (qPCR) nor the level of RlmN protein (targeted proteomics) was affected significantly by erythromycin treatment (**Fig. 3B, C**). These data suggested that the activity of RlmN was regulated post-translationally by antibiotic exposure.

**Figure 3.**
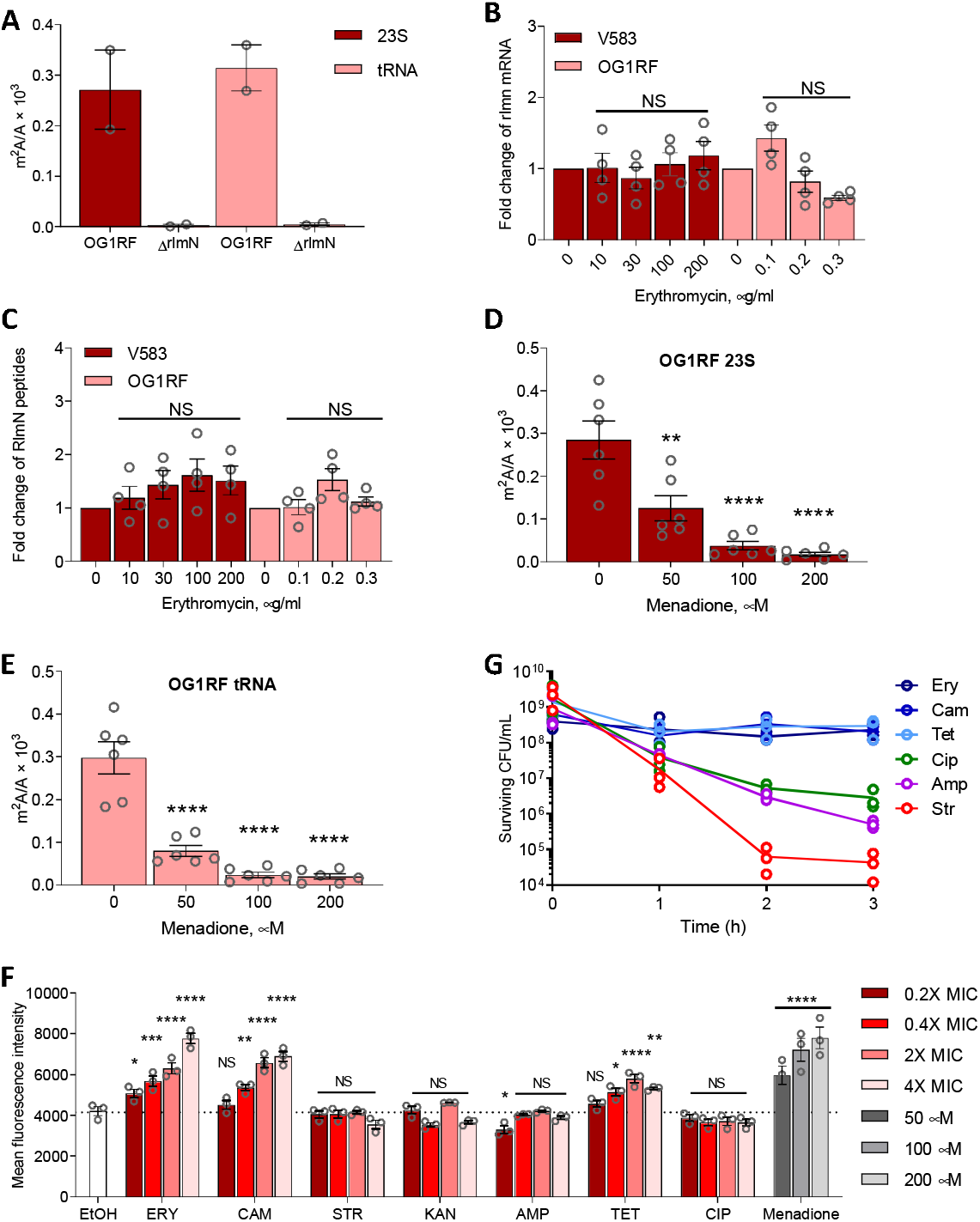
RlmN is regulated at the protein level by reactive oxygen species (ROS). (**A**) Loss of *rlmN* abolishes m^2^A in OG1RF (mean ± deviation about the mean, *n□=□2*). (**B**) RT-qPCR of *rlmN* in OG1RF and V583 upon erythromycin treatment. Data represent mean ±SEM for four experiments: duplicate analyses with two different primer sets. Statistical analysis by one-way analysis of variance (ANOVA) with Dunnett’s test versus untreated: NS, not significant. (**C**) Targeted proteomics of RlmN in OG1RF and V583 upon erythromycin treatment. Data represent mean ± SEM for six experiments: three peptides monitored in two independent experiments. Statistical analysis by one-way analysis of variance (ANOVA) with Dunnett’s test versus untreated: NS, not significant. Ratio of m^2^A to A in (**D**) 23S rRNA and (**E**) tRNA with menadione treatment. Data represent mean ± SEM for three independent experiments performed with technical duplicates. Statistical analysis by one-way analysis of variance (ANOVA) with Dunnett’s test versus untreated: P<0.005 and P<0.0001 are denoted as ** and **** respectively. (**F**) Mean fluorescence intensity of CellROX Green Dye+ in OG1RF treated with various antibiotics at indicated concentrations above and below MICs. Data represent mean ± SEM for three independent experiments. Statistical analysis by twoway analysis of variance (ANOVA) with Dunnett’s test versus EtOH: NS, not significant; P<0.05, P<0.005, P<0.0005 and P<0.0001 are denoted as *, **, *** and **** respectively. (**G**) Cell killing kinetics of various antibiotics at 10X MIC reveal that ROS-generating antibiotics are not bactericidal in OG1RF. Symbols represent individual data points for three independent experiments. Source data are provided as a **Source Data** file.

One possible mechanism for regulating RlmN activity involves antibiotic-induced oxidative stress. RlmN is a radical S-adenosylmethionine (SAM) enzyme containing a [4Fe-4S] cluster that is sensitive to disruption by reactive oxygen species (ROS)^7^. To test this model, we grew OG1RF in the presence of the superoxide radical generator, menadione^8^, at sub-inhibitory concentrations and then measured m^2^A in 23S rRNA and tRNA. Menadione treatment caused a dose-dependent decrease in m^2^A in both rRNA and tRNA (**Fig. 3D-E**). This result implies that the macrolides and chloramphenicol also induced ROS in OG1RF and V583, which we assessed using the fluorogenic superoxide-specific probe CellROX Green to quantify antibiotic-induced ROS levels in OG1RF^9^. As expected, both erythromycin and chloramphenicol increased CellROX Green fluorescence at sub-inhibitory and higher concentrations (**Fig. 3F**), with analysis of forward and side scatterplots suggesting that bacteria morphology remains unchanged, thus ruling out artifacts known to confound the detection of antibiotic-induced ROS^9^ (**Supplementary Fig. 3**). Not all ribosome-targeting antibiotics generate ROS in OG1RF. Sub- and supra-inhibitory concentrations of tetracycline, which binds to the 30S subunit, induced an increase in CellROX Green fluorescence whereas the aminoglycosides streptomycin and kanamycin did not (**Fig. 3F**). Further, the mechanistically distinct ampicillin and ciprofloxacin also did not increase CellROX Green fluorescence at both sub- or supra-inhibitory concentrations (**Fig. 3F**). These observations contradict Léger *et al.*, who reported that the **β**-lactams amoxicillin and cefotaxime caused superoxide production by reduction of demethylmenaquinone (DMK) in *E. faecalis*^10^. However, *E. faecalis* is well established to produce high levels of extracellular superoxide by way of externally-facing, membrane-bound DMK, but the superoxide is unable to diffuse through cell walls and be detected as intracellular superoxide by CellRox and is also rapidly dismutated to hydrogen peroxide outside the cell^11^; Léger *et al.* measured extracellular hydrogen peroxide as their surrogate for superoxide^10^. Along with the menadione data, our results establish that RlmN is inactivated by intracellular superoxide resulting from exposure of *E. faecalis* to macrolides and chloramphenicol.

Given the controversial model that bactericidal antibiotics share a common mechanism of generating cytotoxic ROS^9,12,13^, we tested antibiotics for their bactericidal and bacteriostatic activity in *E. faecalis*. Erythromycin and chloramphenicol have been classified as bacteriostatic^14^, with bactericidal activity at high concentrations against *Streptococcus pneumoniae*^14^, while ciprofloxacin, ampicillin, and streptomycin are classified as bactericidal^14^. However, our data show that erythromycin and chloramphenicol induce superoxide production at low concentrations in OG1RF, while ciprofloxacin, ampicillin, and streptomycin do not. Léger *et al.* observed ROS generation by the related **β**-lactam, amoxicillin, at supra-lethal concentrations, again by measuring hydrogen peroxide which was likely generated from superoxide produced extracellularly. To establish bactericidal activity, we performed killing assays with OG1RF in the presence of antibiotics at ten times their MIC and found that antibiotics that increase CellROX Green fluorescence (*i.e*., erythromycin, chloramphenicol, tetracycline) are bacteriostatic in OG1RF, while ciprofloxacin, ampicillin, and streptomycin, which do not induce ROS, are bactericidal (**Fig. 3G**). These data not only further disprove the link between bactericidal antibiotics, ROS, and cell death, but also raise the question of the role of RlmN in *E. faecalis* antibiotic sensitivity.

**Extended Data Figure 1.**
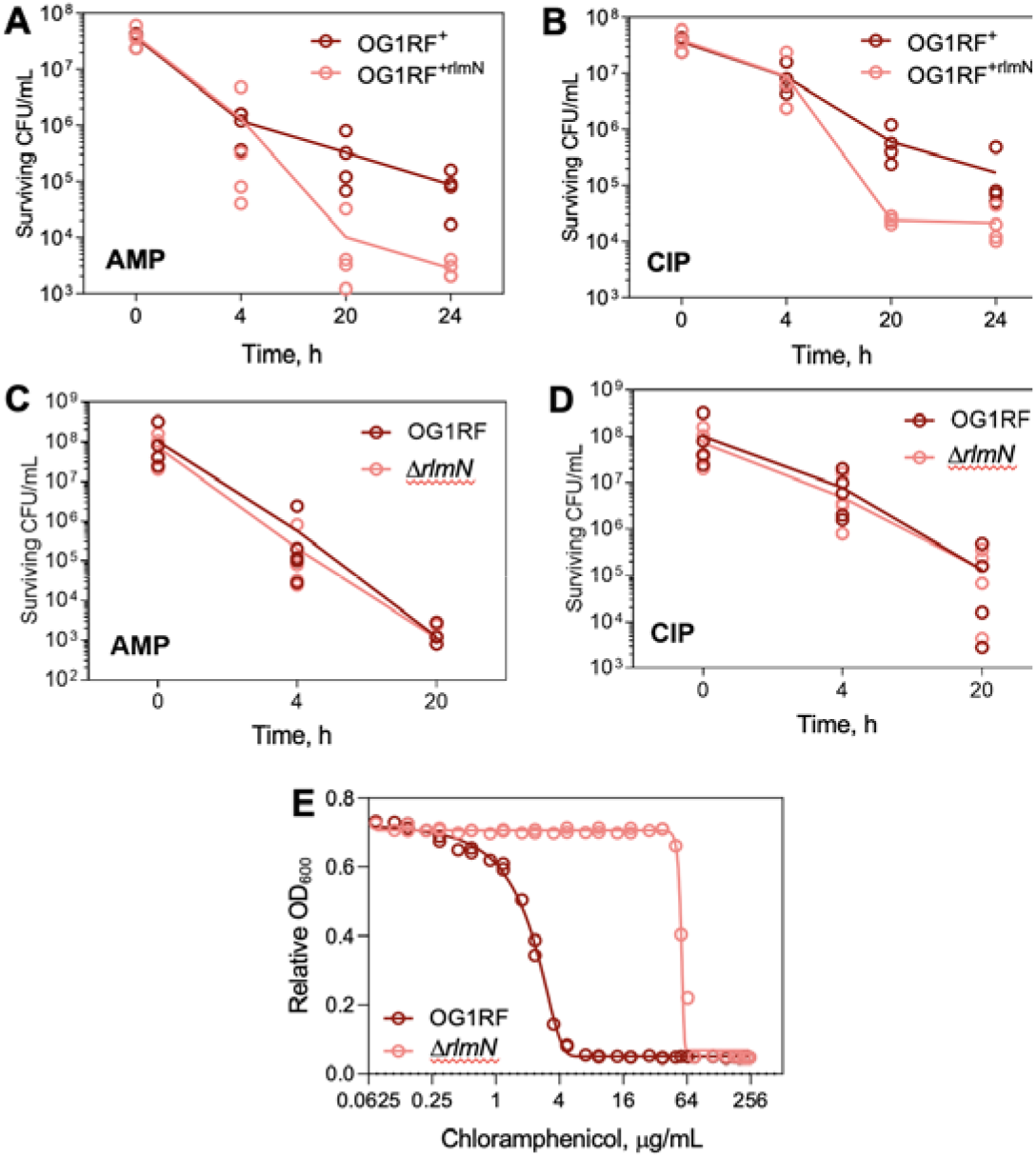
Phenotypic characterization of *rlmN* KO *(ΔrlmN)* and overexpressed RlmN (OG1RFp*rlmN*). Kinetics of cell killing for OG1RFp *Empty* and OG1RFp*rlmN* grown with 5X MIC for (**A**) ampicillin (5 μg/mL) and (**B**) 5 μg/mL ciprofloxacin. Kinetics of cell killing for OG1RF and *ΔrlmN* grown with 5X MIC for (**C**) 10 μg/mL erythromycin and (**D**) 5 μg/mL ciprofloxacin. Graphs show individual data for 4 independent experiments. (**E**) Growth assay for minimum inhibitory concentration (MIC) of chloramphenicol with OG1RF and *ΔrlmN*. The graph shows data

We next investigated the effect of RlmN activity on *E. faecalis* antibiotic sensitivity, using the *ΔrlmN* deletion mutant and a strain over-expressing RlmN. Here we created a constitutive over-expression mutant, OG1RFp*rlmN*, which carried an RlmN-expressing plasmid under the constitutive Sortase A promoter in vector pGCP123^15^; a control strain, OG1RFp*Empty*, carried the same plasmid lacking the coding sequence for RlmN. Over-expression of RlmN did not affect the MIC for erythromycin, streptomycin, kanamycin, ampicillin, tetracycline, or ciprofloxacin (**Supplementary Table 2**), but did increase the sensitivity of OG1*RFprlmN* to killing by the bactericidal antibiotics ampicillin and ciprofloxacin by 10-fold (**Extended Data Fig. 1A, B**). Loss of RlmN however, had no effect on *E. faecalis* growth in the presence of up to 5-times the wild-type MIC for the five antibiotics (**Supplementary Table 2, Extended Data Fig. 1C, D**), except for a 16-fold increase in MIC for chloramphenicol (**Extended Data Fig. 1E, Supplementary Table 3**). Taken together, (1) over-expression of RlmN increases *E. faecalis* sensitivity to ampicillin and ciprofloxacin and (2) loss of RlmN activity confers resistance to chloramphenicol suggest that RlmN may play a role in phenotypic antimicrobial tolerance or resistance.

We next defined the effects of RlmN activity on the cell proteome. Our results establish that RlmN’s methyltransferase activity is attenuated by superoxide, raising the possibility that RlmN serves as a redox sensor that regulates cell stress response. To test this idea, we performed quantitative proteomics using multiplexed isobaric Tandem Mass Tags (TMT) to identify differentially expressed proteins between OG1RF, *ΔrlmN*, and OG1RF grown in menadione (**Fig. 4A, B**; **Supplementary Data 1**). Only a few proteins were upregulated in *ΔrlmN* as compared to OG1RF treated with menadione, suggesting precision of RlmN as a molecular switch. Indeed, we found a positive correlation of R^2^= 0.749 between statistically significant (p<0.05) protein changes in *ΔrlmN* and OG1RF treated with menadione (**Fig. 4D**). One of these was the antioxidant defense enzyme, superoxide dismutase, for which there is a single gene *(sodA)* in *E. faecalis*^16^. Of note, another well-known antioxidant defense enzyme, catalase (KatA), was not consistently detected in the proteomic analyses (**Supplementary Data 1**). Besides SodA, proteins with levels increasing in common (**Fig. 4C**, **Supplementary Table 4**) between *ΔrlmN* and menadione-treated OG1RF include two ribosomal proteins (RpsI, RplE), a DNA repair enzyme LigA, a tRNA-modifying enzyme MnmE, an oxidoreductase, and a ribonucleotide reductase. While there is no obvious gene ontology link among the proteins whose levels increased in *ΔrlmN* (**Supplementary Table 5**), DNA repair, tRNA modification changes, and increases in the dNTP pool are common features of bacterial stress responses^1,17^.

**Figure 4.**
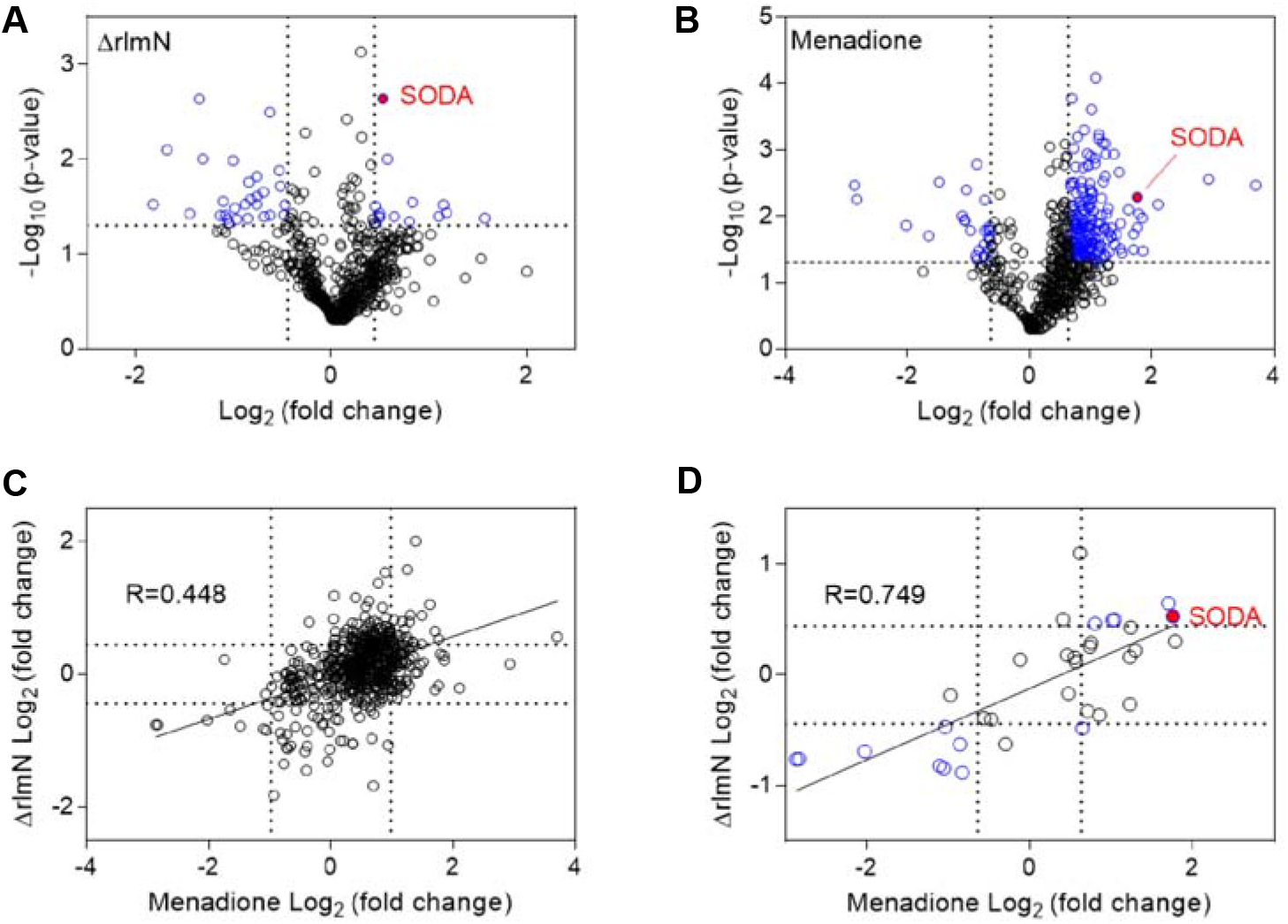
Loss of *rlmN* and treatment with menadione cause similar changes in the OG1RF proteome. Volcano plots showing changes in protein levels in OG1RF caused by (**A**) *rlmN* knockout *(ΔrlmN)* and (**B**) menadione treatment. P-values were calculated using the Student’s t-test. The log_2_ fold-change (x-axis) was plotted against the −log_10_(p value) (y-axis). A −log_10_(p-value) of 1.30 threshold (P<0.05) Is denoted by the dotted horizontal line, while the vertical dotted lines represent fold-change values at ±1 standard deviation from the mean. (**C**) Proteins detected in both menadione treated and *ΔrlmN* without a P-value cutoff. (**D**) Proteins that changed significantly (P<0.05) in both menadione treated and *ΔrlmN* cells. Cut-off was set at 1 standard deviation, which is log_2_(fold change) of ±0.6388 for menadione treatment and log_2_(fold change) of ±0.4432 for *ΔrlmN*. Blue circles represent proteins which significantly up- or down-regulated relative to untreated, wild-type cells. These proteins are shown in **Supplementary Table 4**. Data shown are of three biological replicates. Red circles denote superoxide dismutase (SODA).

Interestingly, the common set of proteins which are downregulated in both datasets include proteins associated with virulence. Pilus subunit proteins EbpA and EbpB are major virulence factors in *E. faecalis*, involved in biofilm formation, endocarditis, and catheter-associated urinary tract infection^18^. Phosphocarrier protein HPr, a component of the phosphoenolpyruvate-dependent sugar phosphotransferase system, has been found to contribute to successful bacteremia of *Neisseria meningitidis* in a mice infection model^19^, and activation of virulence in *Listeria monocytogenes*^20^. WxL domain-containing proteins in *Enterococus faecium* have been implicated in survival in bile salt and the pathogenesis of endocarditis^21^. Other proteins include a signal peptidase and a CAAX amino protease family protein, of which the latter is predicted to be involved in regulation of transcription.

Based on the results presented here, we propose a mechanism (**Extended Data Fig. 2**) in which RlmN serves as a ROS-sensitive molecular switch that modulates physiological responses to oxidative stress and confers phenotypic resistance to environmental stresses and antimicrobial agents. This is perhaps not surprising given the importance of other Fe-S proteins, such as SoxR, Fnr, and aconitase, as ROS sensors linked to changes in gene expression and cell phenotype^22^. RlmN and m^2^A are absent in eukaryotes and, within prokaryotes, RlmN is the only enzyme that synthesizes m^2^A. The inertness of the C2 of adenosine to electrophilic attack and the low acidity of the C2 proton requires a free radical SAM intermediate unique to RlmN^23^ and the chloramphenicol-florfenicol resistance methyltransferase (Cfr)^24^. Here we showed that RlmN activity is not only strongly dependent upon intracellular superoxide levels but also regulates levels of SodA, which promotes antibiotic tolerance^25^ and facilitates survival in macrophages^16^ in *E. faecalis* and other bacteria.

**Extended Data Figure 2.**
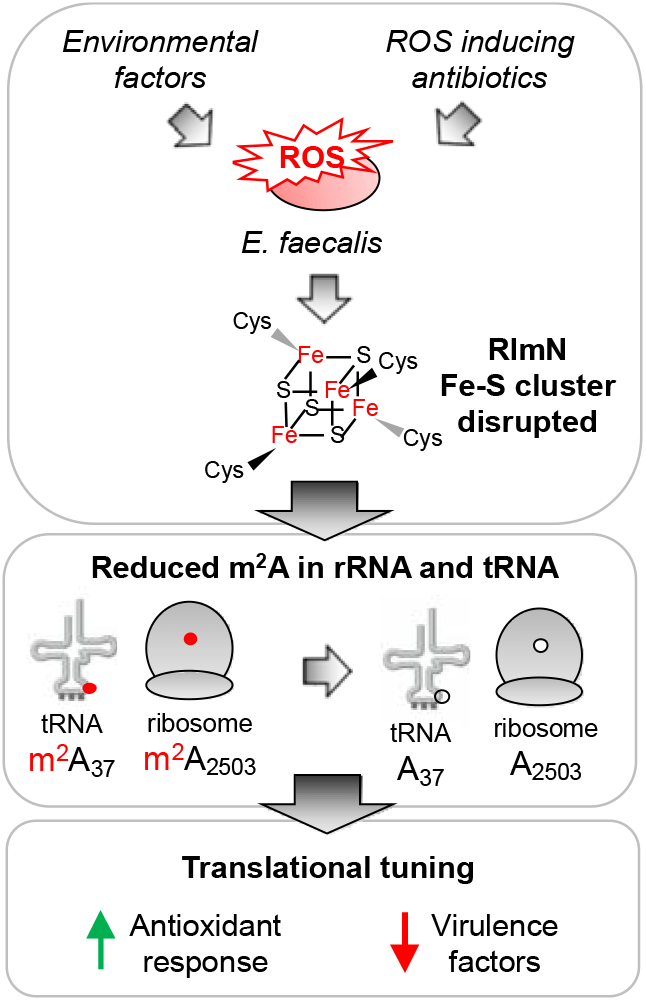
Working model of RlmN as a sensor for oxidative stress. Environmental factors as well as certain antibiotics induce reactive oxygen species in *Enterococcus faecalis*. These lead to the inactivation of RlmN through perturbation of its Fe-S cluster, exerting dual effects on both the ribosome and tRNA. m^2^A is present in position 37 of selected tRNAs and 23S rRNA in the peptidyl transferase center of the ribosome. Chemical structure of m^2^A with the 2-methyl group shown in red. Through loss of modifications on its rRNA and tRNA, and lead to a modified pool of ribosomes and tRNA, which causes the upregulation of SOD for improved survival in the presence of antibiotics.

So how does exposure of *E. faecalis* to ribosome-binding antibiotics lead to elevated superoxide levels? While there is no universal mechanism by which antibiotic exposure causes increases in ROS in bacteria, in spite of earlier claims^13^, there are numerous pathways for generating superoxide and other ROS and for environmental exposure of bacteria to ROS^7^. *E. faecalis* generates large amounts of extracellular superoxide^11^, with Léger *et al.* showing that supra-lethal ampicillin doses increase these levels^10^. While we did not measure extracellular ROS, superoxide cannot diffuse through the bacterial cell wall and our results show that neither sub-nor supra-lethal amoxicillin or other bactericidal antibiotics cause intracellular superoxide formation in *E. faecalis* (**Fig. 3f**). How sublethal concentrations of erythromycin and chloramphenicol cause superoxide levels to increase could relate to the cell stress caused by inhibition of translation or by mistranslation, with the proteotoxic stress leading to increased reductive metabolism and thus increases in superoxide. The mechanism of erythromycin-induced superoxide production awaits further study.

How does RlmN-catalyzed m^2^A play a role in the phenotypic changes caused by loss of RlmN or exposure to superoxide? Two possibilities come to mind based on RlmN activity on both rRNA and tRNA. From the rRNA perspective, m^2^A might facilitate the ribosome stalling that leads to ErmBL nascent peptide activation of ErmB expression^26^. RlmN catalyzes m^2^A formation at A2503 of 23S rRNA, a conserved nucleotide which resides in the peptidyltransferase center (PTC) of the ribosome near the entrance to the exit channel for the nascent polypeptide (**Extended Data Fig. 3**) and is involved in fine-tuning ribosome–nascent peptide interactions, relaying the stalling signal to the PTC^27^. A2503 is very close spatially to the erythromycin binding site^27^, to the ErmBL nascent peptide that activates ErmB expression upon antibiotic binding^28^, and to the A2058 that is modified with m^6^A by ErmB to prevent antibiotic binding (**Extended Data Fig. 3**). Though speculative, one hypothesis is that loss of RlmN activity reduces m^2^A2503 and thus facilitates ErmBL-induced activation of ErmB synthesis.

**Extended Data Figure 3.**
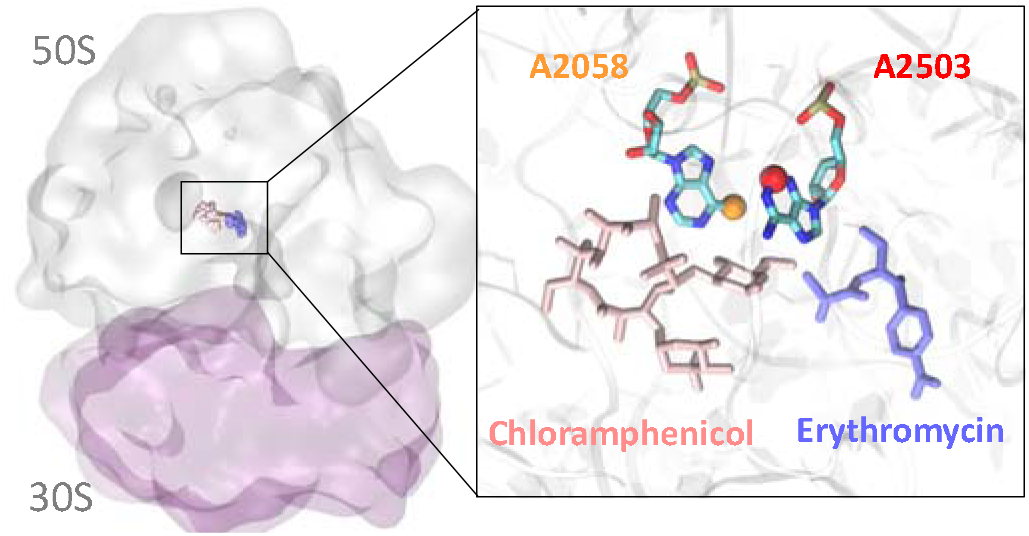
Structure of the ribosome showing superimposed binding sites for erythromycin (blue), chloramphenicol (pink), the A2503 (red ball) modified with m^2^A by RlmN, and the A2058 site (yellow ball) modified with Erm-mediated m^6^A.

From the tRNA perspective, RlmN is one of only two methyltransferases known to modify both rRNA and tRNA^3^. In *E. coli*, m^2^A37 occurs in six tRNAs: tRNA^Arg^ICG, tRNA^Asp^QUC, tRNA^Gln^cmnm^5^sUUG, tRNA^Gln^CUG, tRNA^Glu^mnm^5^s^2^UUC, and tRNA^His^QUG^29^. Modifications at position 37 are important for maintaining the reading frame^30^, while loss of RlmN increases stop codon readthrough^31^. The latter is likely not relevant for synthesis of selenoproteins in *E. faecalis* since there are no apparent genes encoding selenoproteins in the *E. faecalis* genome.^32^ Further studies are underway to determine if *E. faecalis* uses tRNA reprogramming and codon-biased translation to regulate expression of stress response genes as observed in mycobacteria^1^.

Finally, our results raise the question of RlmN sensitivity to inactivation by superoxide: is RlmN unique in its sensitivity compared to other Fe-S cluster-containing proteins in *E. faecalis*? *E. faecalis* is unusual among human commensal and pathogenic bacteria in lacking many Fe-S cluster proteins, such as fumarase, aconitase, isocitrate dehydrogenase, and succinate dehydrogenase in the tricarboxylic acid cycle^33^. This lack of a tricarboxylate cycle is shared by *Listeria monocytogenes*, in which bactericidal antibiotics have also been shown not to produce ROS^34^. Other Fe-S cluster proteins absent in *E. faecalis* include the GrxD iron transport regulator, 2 of 3 systems for Fe-S cluster biogenesis (NIF, ISC, and SUF; only SUF is present in *E. faecalis*^35^), and MiaB. The latter is corroborated by our inability to detect m^2^si^6^A in the presence of i^6^A (**Fig. 1D**). However, *E. faecalis* possesses other Fe-S cluster proteins, including QueE and QueG involved in queuosine (Q) biosynthesis^36^. Clearly more work is needed to determine the effect of superoxide and other ROS on the activities of different Fe-S cluster proteins to determine if RlmN is uniquely sensitive as a potential redox signaling node in *E. faecalis*.

In conclusion, we showed that RlmN activity is not only strongly dependent upon superoxide levels but also regulates levels of SodA. In *E. faecalis* and other bacteria, SodA promotes antibiotic tolerance^25^ and facilitates survival in macrophages^16^. In all, RlmN, widely distributed across bacteria genera,^37^ may serve as a redox switch relaying redox sensing to both the rRNA and tRNA epitranscriptome for direct modulation of translation for protective oxidative stress response.

## Methods

### Bacteria strains, plasmids, and growth conditions

*Enterococcus faecalis* strains OG1RF and V583 are grown in tryptic soy broth (TSB) or plated on tryptic soy agar under aerobic conditions at 37 ^o^C. All mutant strains are derivatives of OG1RF. The *rlmN* knock-out in OG1RF *(ΔrlmN)* was generated by an in-frame deletion of *rlmN* from OG1RF by allelic replacement using vector pGCP213^15^. RlmN over expressor, OG1RFp*rlmN* was generated by the introduction of the gene coding for RlmN into the plasmid pGCP123 under the constitutive sortase promoter^15^. OG1RFp*Emptf* is OG1RF carrying the plasmid pGCP123, but without introduction of the gene coding for RlmN. All the plasmids used in this study are listed in **Supplementary Table 6**. All the oligos for making the mutant strains are listed in **Supplementary Table 7**. All plasmid constructions are described in **Supplementary Methods** and were verified by Sanger sequencing. MICs used for OG1RF are 1 μg/mL erythromycin, 1 μg/mL spiramycin, 4 μg/mL chloramphenicol, 64 μg/mL kanamycin, 128 μg/mL streptomycin, 1 μg/mL ampicillin, 0.5 μg/mL tetracycline and 1 μg/mL ciprofloxacin. MICs used for V583 are >1024 μg/mL erythromycin, 8 μg/mL chloramphenicol, 256 μg/mL gentamicin, 1 μg/mL ampicillin and1 μg/mL ciprofloxacin. OG1RFp*rlmN* and OG1RFp*Empty* are grown in 500 μg/mL kanamycin to maintain the pGCP123 plasmid.

### Identification and quantification of RNA modifications

An overnight culture was diluted 1:20 fold and then grown at 37 °C to reach mid-log phase. This mid-log phase culture was then diluted 1:20 into media containing sub-lethal concentrations of antibiotics and grown at 37 °C with shaking at 180 rpm. The cultures are harvested at an optical density (OD_600_) of ~0.6-0.8 after 4-5 doublings by centrifugation at 4,000 xg for 10 min at 4 ^o^C. RNA was extracted and purified following Hia *et al.*^38^ Briefly, 100 mL of bacteria culture was lysed in the presence of phenol:chloroform:isoamyl alcohol and 100 mM sodium acetate pH 5.0 by bead beating with 0.1-mm zirconia-silica beads using Qiagen TissueLyser II for 12 min at 30Hz. Large and small RNA species were differentially recovered using the PureLink miRNA Isolation Kit (Invitrogen) with 35% ethanol and 70% ethanol respectively. 23S and 16S rRNA are separately isolated to purity from the large RNA fraction following HPLC using the Bio SEC-5 column (Agilent; 7.8 mm, length: 300 mm, particle size: 5 μm, pore size: 1000 Å); and tRNA was isolated to purity from the small RNA fraction following HPLC on the Bio SEC-3 column (Agilent; 7.8 mm, length: 300 mm, particle size: 5 μm, pore size: 300 Å). All separations were run with 100 mM ammonium acetate, at 60 °C with a flow rate of 0.5 mL/min.

Peaks corresponding to the right RNA populations are concentrated and subjected to buffer exchange into 10 mM Ammonium Acetate using size exclusion filters (Millipore). RNA (5 μg) was enzymatically hydrolyzed for 4 h at 37 °C with benzonase (99% purity, Novagen 70664), bacterial phosphatase (ThermoFisher 18011015) and phosphodiesterase I (Sigma P3243), in the presence of magnesium chloride, antioxidants and deaminase inhibitors including desferroxamine (Sigma D9533), butylated hydroxyltoluene (Sigma W218405), pentostatin (Sigma SML0508), tetrahydrouridine (Calbiochem 584222), and internal standard [^15^N]_5_-deoxyadenosine. Samples were cleaned up using a 10 kDa cut-off filter (Nanosep).

The ribonucleoside mixtures were separated on a Hypersil C18 analytical column (2.1x 100 mm, 1.8 mm; Agilent) on an Agilent 6490 QQQ triple-quadrupole LC mass spectrometer using multiple reaction monitoring in positive-ion mode as previously described^1^. The nucleosides were identified using the retention time of the pure standards and the nucleoside to base ion mass transitions involving loss of either ribose (136 *m/z)* or 2’-*O*-methyl-ribose (146 *m/z)* (**Supplementary Table 1**). For relative quantitation of modifications among the same batch of samples, the signal intensity is normalized against the combined intensity of the four canonical ribonucleosides to correct for variation in RNA quantities. Spectral signals are also normalized against spiked internal standard ([^15^N]_5_-2’-deoxyadenosine) to adjust for variations in instrument sensitivity. For absolute quantification of m^2^A and adenosine, a series of concentrations of nucleoside standards for m^2^A and adenosine were run with every batch of samples to obtain standard calibration curves. The concentrations of nucleosides were then obtained by fitting the signal intensities onto the calibration curves, and these were then used to obtain the molar ratio of m^2^A/A.

### Measurement of *rlmN* mRNA levels

Total RNA (2 μg) was subjected to DNA removal using the TURBO DNA-free kit (Ambion, Life Technologies) following manufacturer’s protocol. 600 ng in 15 μL was used for reverse transcription using the iScript cDNA synthesis kit (Bio-Rad, Hercules, CA, USA). The reverse transcription program was run as follows: 25 °C for 5 min, 42 °C for 30 min and 85 °C for 5 min, followed by a cooling step at 4 °C. Two-step real-time quantitative polymerase chain reaction (qPCR) was then performed using the BlitzAmp qPCR mastermix (MiRxes, Singapore). Primer sequences can be found in **Supplementary Table 8**. The qPCR program was run as follows: 95 °C for 5 min followed by 40 cycles of denaturation at 95 °C for 10 s and annealing/extension at 60 °C for 30 s. A melting curve analysis consisting of 0.5 °C increments from 65 to 95 °C was performed for all reactions to ascertain the specificity of the primers. RpoA served as internal loading control.

### Measurement of RlmN protein levels

Bacteria pellets from 10 mL of log phase culture of OG1RF and V583 were resuspended in 250 μL of 50 mM Hepes pH 8, 8 M urea, 1 mM DTT and homogenized by bead-beating followed by clarification by centrifugation at 16,000 xg for 30 min at 4 °C. Following quantification of total protein in the supernatant using the bicinchoninic acid protein assay (ThermoFisher Scientific), 50 μg of protein was mixed with SDS-PAGE loading dye and separated on a 14% SDS-PAGE gel. Following staining and destaining, a gel slice corresponding to 30-45 kDa, encompassing target protein RlmN 40.9 kDa and reference proteins RpoA 35.05 kda and Gap2 35.77 kDa, was excised and cut into 1-2 mm pieces. The gel pieces were destained, reduced and alkylated followed by overnight trypsin digestion and peptide extraction following manufacturer’s instructions (In-Gel Tryptic Digestion kit, ThermoFisher Scientific). Extracted peptides were vacuum dried and redissolved in 2% acetonitrile in 0.1% formic acid in water.

Skyline (http://proteome.gs.washington.edu/software/skyline) was used to identify precursor peptides and transitions to be used for the targeted quantification of RlmN and reference proteins (**Supplementary Table 3**). Automated picking of precursor peptides and transitions was used with filters set to select for singly charged, long (y3 and greater) y-ions with no *m/z* overlaps with b-ions. Selected peptides were synthesized at 90% purity and analyzed on a Hypersil C18 analytical column (2.1x 100 mm, 1.8 mm; Agilent) on an Agilent 1290 infinity LC system coupled to an Agilent 6490 QQQ spectrometer in positive ion mode. Agilent Automated MRM Method Optimizer for Peptides was used to optimize collision energies and fragmentation voltages for their MRM transitions and peptides were used at a concentration of 10 μg/mL for determination of retention times.

Reversed-phase chromatography was performed with a fixed flow rate of 0.25mL/min with a gradient of water and acetonitrile (solvent B) acidified with 0.1% (v/v) formic acid. Gradients used were as follow: 0-29% solvent B from 0-29 min, 29-90% from 29-30 min, 90% for 38 min, 90% to 0 % from 38-39 min, and 0% for 45 min. Source conditions: gas temperature 325 °C, gas flow 10 L/min, nebulizer 32 psi, sheath gas temperature 300 °C, sheath gas flow 11 L/min, capillary 2000 V, charging 500 V. Columns were incubated at 40 °C. The top two precursor peptides by peak area and number of transitions (minimum 2) were selected as qualification and quantification ions.

### Flow cytometry assays

CellROX Green (ThermoFisher) is a proprietary oxidation-sensitive dye whose fluorescence at 500–550 nm after excitation at 488 nm increases substantially on oxidation in the presence of dsDNA. Cellrox green reacts to hydroxyl radical and superoxide but not hydrogen peroxide^9^. Log-phase cultures were diluted to OD_600_ of 0.1 in 10% TSB in the presence of menadione or antibiotics and incubated for 30 min at 37 °C followed by the addition of CellROX green (final 0.5 μM) for a further 30mins at 37 °C in the dark with shaking at 180 rpm. Samples were analyzed using a HTS fluidics system and the flow rate was set to 3.0 mL/s with a 150 mL injection volume, 100 mL mixing volume, 250 mL/s mixing speed, five mixes, and a wash volume of 800 mL. Samples were analyzed on a custom LSR II flow cytometer (BD Biosciences), detected with a 530/30 nm band-pass emission and recording 50,000 events. Data were analyzed with FlowJo v10.0.6 (Tree Star, Inc.). Gating was set using unstained samples for the bacterial population by forward-scatter (FSC; correlates with cell size) and side-scatter (SSC; correlates with cell internal granularity) of light and to determine background fluorescence.

### Proteomics

Fresh mid-log phase cultures (OD_600_ of 0.6) were diluted 1:20 into TSB media with or without sub-lethal concentrations of antibiotics or menadione and grown at 37 °C with shaking at 180 rpm. The cultures are harvested at an optical density (OD_600_) of ~0.6-0.8 after 4-5 doublings by centrifugation at 4,000 xg for 10 min at 4 °C. Bacterial pellets were resuspended in 250 μL of 50 mM Hepes pH 8, 8 M urea, 1 mM DTT, and homogenized by bead-beating followed by clarification by centrifugation at 16,000 xg for 30 min at 4 °C. Following protein quantification in the supernatant using the bicinchoninic acid protein assay (ThermoFisher Scientific), 200 μg of protein was digested with trypsin after being reduced with DTT and alkylated with iodoacetamide. After washing 3 times with 0.5 M TEAB followed by fractionation using the 10 kDa ultrafiltration system, ~100 μg of digested peptides from each group, including two biological replicates, was labeled using the six-plex TMT isobaric and isotopic mass-tagging kit (ThermoFisher Scientific), which was performed according to manufacturer’s instructions.

Peptides were separated by reverse phase HPLC (Thermo Easy nLC1000) using a precolumn (made in house, 6 cm of 10 μm C18) and a self-pack 5 μm tip analytical column (15 cm of 5 μm C18, New Objective) over a 150 min gradient before nanoelectrospray using a QExactive mass spectrometer (ThermoFisher). The mass spectrometer was operated in a data-dependent mode. The parameters for the full scan MS were as follow: resolution of 70,000 across 350-2000 *m/z*, AGC 3e6, and maximum IT 50 ms. The full MS scan was followed by MS/MS for the top 15 precursor ions in each cycle with a NCE of 28 and dynamic exclusion of 30 s. Raw mass spectral data files (.raw) were searched using Proteome Discoverer (ThermoFisher) and Sequest (Reference). Search parameters were as follow: 10 ppm mass tolerance for precursor ions; 0.8Da for fragment ion mass tolerance; 2 missed cleavages of trypsin; fixed modification was carbamidomethylation of cysteine, and N-term and Lysine TMT-label; variable modifications were methionine oxidation and serine, threonine and tyrosine phosphorylation. Only peptides with a Scorer score ≥2, reporter channel intensity >500 and an isolation interference ≤30 were included in the data analysis. Proteomics data are presented in **Supplementary Data 1**.

### Bactericidal activity analysis

A log phase culture of OG1RF was diluted to a final concentration of 10^8^ CFU/mL (OD_600_ ~0.1) in Tryptic Soy Broth in the presence of antibiotics at 37 °C with shaking. Aliquots were drawn from each respective tube at various time points and serially diluted until 10^-8^-fold. An aliquot (2.5 μL) of each dilution was spotted onto TSB agar and incubated at 37 °C overnight, with colonies counted 24 h after spotting.

### Determination of minimal inhibitory concentrations (MIC) of antibiotics

Two-fold serial dilutions of antibiotics in TSB were performed in separate rows of a polystyrene 96-well plate (Corning) with each plate containing an inoculum of respective bacteria. The inoculum was a 1:500 dilution from a culture at log phase (OD_600_ = 0.5) grown at 37 °C. The plate was incubated with shaking at 37 °C and the optical density of each well was measured at a wavelength of 600 nm (BIOTEK, Synergy 4). The MIC values were taken as the lowest concentration for which no growth was discernible (<0.05 OD_600_) after 24 h. All tests were performed three times independently with two samples in each test. MIC data are presented in **Supplementary Table 2**.

### Antibiotic killing assays

The OG1RF strains were cultured aerobically in TSB 37□°C for approximately 16□h with shaking (180□rpm) followed by 1:20 dilution and cultured to mid-log growth. Strains OG1RFp*Empty* and OG1RFp*rlmN* harboring pGCP123 plasmids were grown in 500 μg/mL kanamycin. The mid-log culture was diluted to OD_600_ ~0.06 (~5×10^7^□cfu/mL). An aliquot was plated to enumerate the colony-forming units (cfu) (Time 0) before the addition of antibiotics with final concentrations at 20 μg/mL ciprofloxacin and 20 μg/mL ampicillin. An aliquot was removed at the indicated time points and washed with sterile PBS. The cells were serially diluted and plated on tryptic soy agar to enumerate the survivors.

### Statistical analysis

Statistical significance was assessed using appropriate tests using Prism 8 (GraphPad) software, detailed in their respective figure legends. Asterisks indicate the level of statistical significance: *P□<□0.05 **P□<□0.01, ***P□<□0.001 and ****P□<□0.0001. P values □ < □ 0.05 were considered significant. Experiments were repeated at least three times.

## Data availability

The raw mass spectrometry data are available at Chorus (chorusproject.org) under accession number 1650. The source data used to generate plots are provided as a **Source Data** file. All other data are available from the corresponding author on request.

## Acknowledgements

The authors thank Dr. Michael DeMott for his assistance with the proteomics work. This research was supported by the National Research Foundation of Singapore through the Singapore–MIT Alliance for Research and Technology Antimicrobial Resistance Interdisciplinary Research Group. Contributions of LNL were supported by the National Research Foundation and Ministry of Education Singapore under its Research Centre of Excellence Programme at SCELSE. AS and JL acknowledge support from the Singapore MIT Alliance (SMA) Graduate Fellowship and MOE Tier 2 Grant MOE2018-T2-2-13. Proteomics work was performed in part in the MIT Center for Environmental Health Sciences Bioanalytical Core, which is supported by Center grant P30 ES002109 from the National Institute of Environmental Health Sciences with the aid of Dr. Michael Demott.

## Author contributions

WLL, KK and PD designed research; WLL, AS, LNL, HLL, JL, PH, LC and PD performed research; CSCC contributed new reagents/analytic tools; WLL, AS, KK, PD analyzed data, all authors participated in writing the manuscript.

The authors declare no competing interest.

## Conflict of interest

The authors declare no conflicts of interest.

## Notes

### Competing Interest Statement

The authors have declared no competing interest.

