## Supplementary Information for "An RNA modification enzyme directly senses reactive oxygen species for translational regulation in *Enterococcus faecalis*"

**Table of contents**

- **Supplementary figures**
  - **Supplementary Figure 1**. LC‐MS/MS extracted ion chromatograms of modified ribonucleosides analyzed in V583 23S and 16S rRNA and tRNA.
  - **Supplementary Figure 2.** Epitranscriptomic profiling of 23S and tRNA of OG1RF grown in the presence of sub-inhibitory concentrations of erythromycin.
  - **Supplementary Figure 3**. Representative contour plots of CellROX Green-stained OG1RF treated with various antibiotics at indicated concentrations.
- **Supplementary tables**
  - **Supplementary Table 1**. Table of ribonucleoside to base ion mass transitions and transition times for monitored ribonucleoside modifications in V583.
  - **Supplementary Table 2.** Minimum inhibitory concentrations (MIC, µg/mL) of V583 and OG1RF WT and strains using a broth microdilution assay
  - **Supplementary Table 3.** Peptides used for targeted protein mass spectrometry
  - **Supplementary Table 4.** Proteins significantly up- and down-regulated in both the Δ*rlmN* mutant following menadione treatment.
  - **Supplementary Table 5.** Gene ontology of molecular function of proteins significantly up-regulated in Δ*rlmN* mutant, as computed by ShinyGO v0.61.
  - **Supplementary Table 6.** Plasmids used in this study
  - **Supplementary Table 7.** Cloning primers used in this study
  - **Supplementary Table 8.** Primers used for RT-qPCR
- **Supplementary methods**

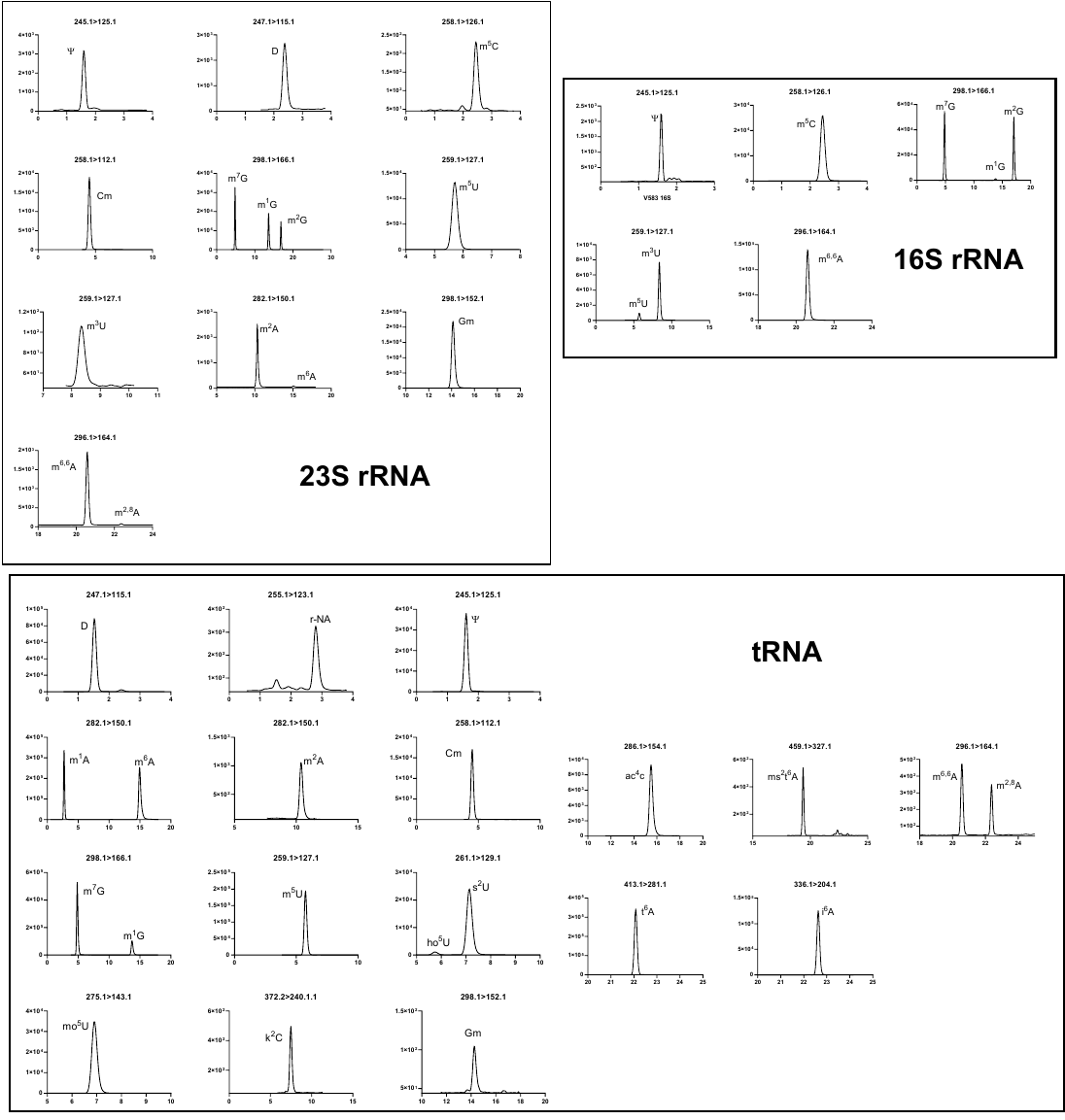

**Supplementary Figure 1.** LC‐MS/MS extracted ion chromatograms of modified ribonucleosides analyzed in V583 23S and 16S rRNA and tRNA. The “X > Y” values denote collision-induced dissociation mass transitions involving the loss of either ribose (136 *m/z*) or 2’-*O*-methyl-ribose (146 *m/z*).

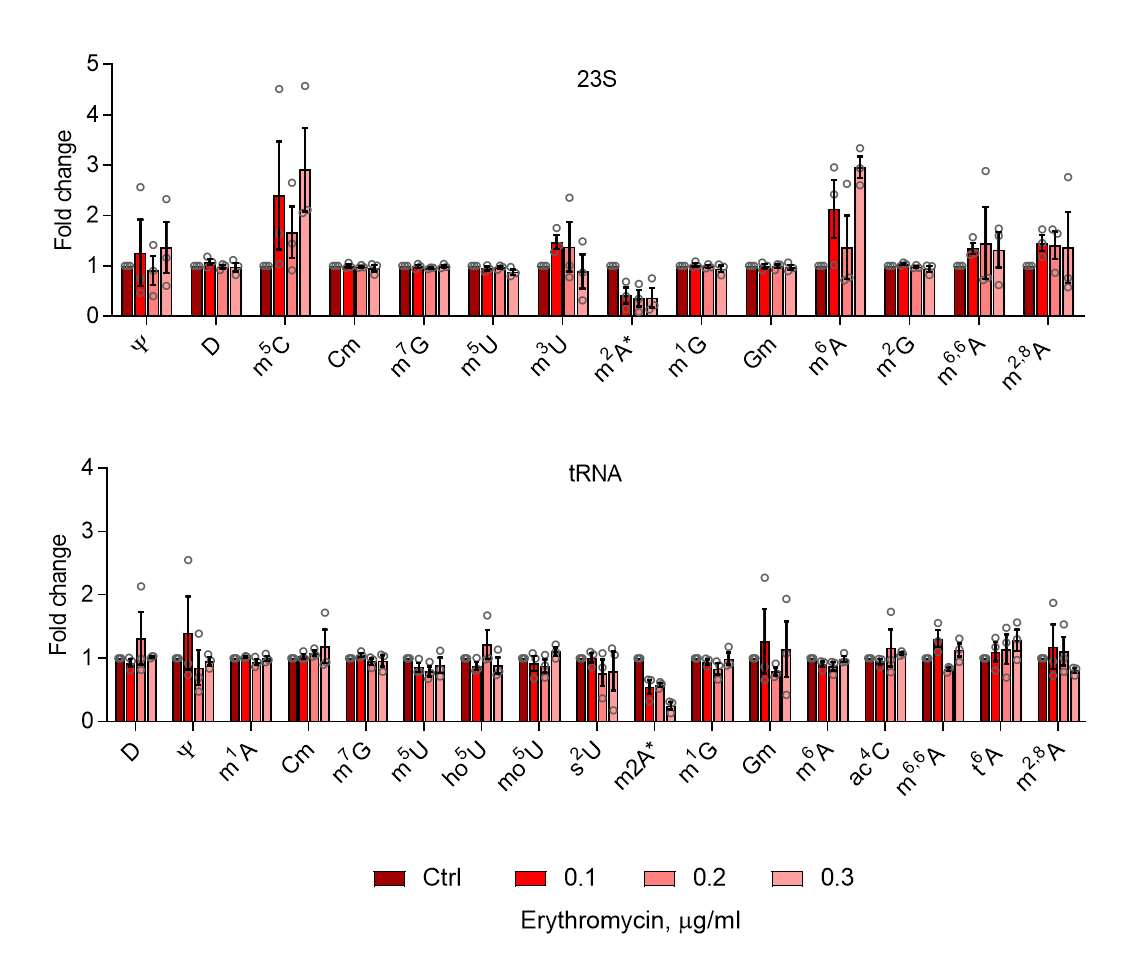

**Supplementary Fig. 2.** Epitranscriptomic profiling of 23S and tRNA of OG1RF grown in the presence of sub-inhibitory concentrations of erythromycin. Changes in RNA modifications in OG1RF in erythromycin (0.1, 0.2 and 0.3 µg/ml) compared to untreated. Levels of modifications are shown as fold change to that within the untreated control. 23S rRNA (top panel) and tRNA (bottom panel). Modifications are arranged from left to right in ascending retention times. m2A is shown with an asterisk. Full names, precursor and product ion masses and retention times of RNA modifications can be found in **Supplementary Table 1**. All data are derived from three independent experiments (mean ± sem, *n* = 3). Source data are provided as a Source Data file.

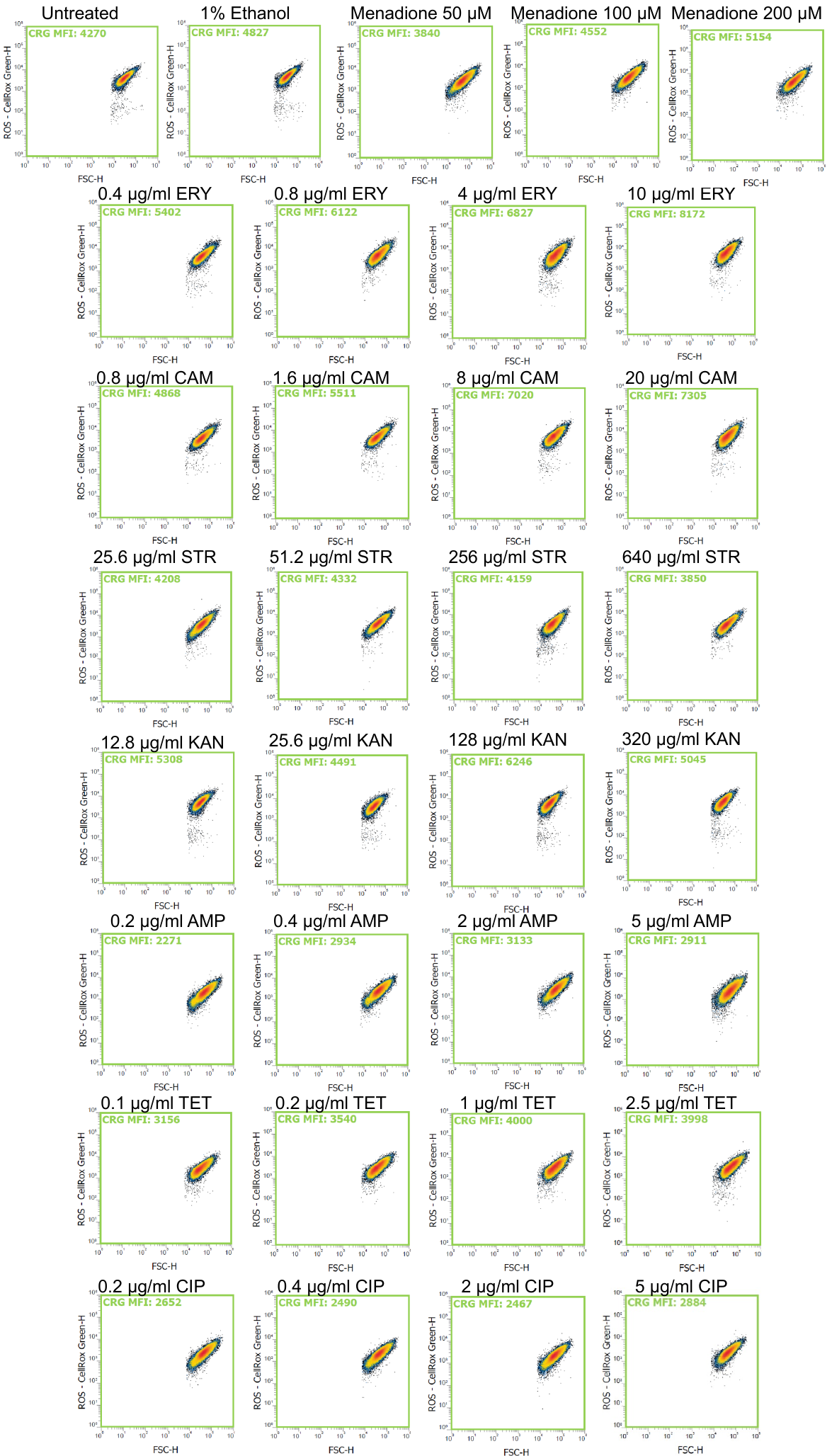

**Supplementary Figure 3**. Representative contour plots of CellROX Green-stained OG1RF treated with various antibiotics at indicated concentrations. ERY, erythromycin, CAM chloramphenicol, STR streptomycin, KAN kanamycin, AMP ampicillin, TET tetracycline, CIP ciprofloxacin.

**Supplementary Table 1.** Table of ribonucleoside to base ion mass transitions and transition times for monitored ribonucleoside modifications in V583. Related to Figure S1.

|  | **Modified ribonucleosides** | **Abbrev** | **Retention Time (min)** | **Precursor ion (*m/z*)** | **Product ion (*m/z*)** | **Neutral Loss (amu)** |
| --- | --- | --- | --- | --- | --- | --- |
| **23S ribonucleosides** | | | | | | |
| 1 | Pseudouridine | Ψ | 1.6 | 245.1 | 125.1 | 120 |
| 2 | Dihydrouridine | D | 2.3 | 247.1 | 115.1 | 132 |
| 3 | 5-Methylcytidine | m^5^C | 2.4 | 258.1 | 126.1 | 132 |
| 4 | 2'-O-Methylcytidine | Cm | 4.5 | 258.1 | 112.1 | 146 |
| 5 | 7-Methylguanosine | m^7^G | 4.8 | 298.1 | 166.1 | 132 |
| 6 | 5-Methyluridine | m^5^U | 5.7 | 259.1 | 127.1 | 132 |
| 7 | 3-Methyluridine | m^3^U | 8.3 | 259.1 | 127.1 | 132 |
| 8 | 2-Methyladenosine | m^2^A | 10.5 | 282.1 | 150.1 | 132 |
| 9 | 1-Methylguanosine | m^1^G | 13.7 | 298.1 | 166.1 | 132 |
| 10 | 2'-O-Methylguanosine | Gm | 14.3 | 298.1 | 152.1 | 146 |
| 11 | 6-Methyladenosine | m^6^A | 15.1 | 282.1 | 150.1 | 132 |
| 12 | 2-Methylguanosine | m^2^G | 16.9 | 298.1 | 166.1 | 132 |
| 13 | N^6^, N^6^-Dimethyladenosine | m^6,6^A | 20.6 | 296.1 | 164.1 | 132 |
| 14 | 2,8-Dimethyladenosine | m^2,8^A | 22.3 | 296.1 | 164.1 | 132 |
| **16S ribonucleosides** | | | | | | |
| 1 | Pseudouridine | Ψ | 1.6 | 245.1 | 125.1 | 120 |
| 2 | 5-methylcytidine | m^5^C | 2.4 | 258.1 | 126.1 | 132 |
| 3 | 7-Methylguanosine | m^7^G | 4.8 | 298.1 | 166.1 | 132 |
| 4 | 5-Methyluridine | m^5^U | 5.7 | 259.1 | 127.1 | 132 |
| 5 | 3-Methyluridine | m^3^U | 8.3 | 259.1 | 127.1 | 132 |
| 6 | 1-Methylguanosine | m^1^G | 13.7 | 298.1 | 166.1 | 132 |
| 7 | 2-Methylguanosine | m^2^G | 16.9 | 298.1 | 166.1 | 132 |
| 8 | N^6^, N^6^-Dimethyladenosine | m^6,6^A | 20.6 | 296.1 | 164.1 | 132 |
| **tRNA ribonucleosides** | | | | | | |
| 1 | Dihydrouridine | D | 1.5 | 247.1 | 115.1 | 132 |
| 2 | N-Ribosylnicotinamide | r-NA | 2.8 | 255.1 | 123.1 | 132 |
| 3 | Pseudouridine | Ψ | 1.6 | 245.1 | 125.1 | 120 |
| 4 | 1-Methyladenosine | m^1^A | 2.7 | 282.1 | 150.1 | 132 |
| 5 | 2'-O-Methylcytidine | Cm | 4.5 | 258.1 | 112.1 | 146 |
| 6 | 7-Methylguanosine | m^7^G | 4.9 | 298.1 | 166.1 | 132 |
| 7 | 3-Methyluridine | m^5^U | 5.8 | 259.1 | 127.1 | 132 |
| 8 | 5-hydroxyuridine | ho^5^U | 5.8 | 261.1 | 129.1 | 132 |
| 9 | 5-methoxyuridine | mo^5^U | 7.0 | 275.1 | 143.1 | 132 |
| 10 | 2-Thiouridine | s^2^U | 7.1 | 261.1 | 129.1 | 132 |
| 11 | 2-Lysidine | k^2^C | 7.5 | 372.2 | 240.1 | 132.1 |
| 12 | 2-Methyladenosine | m^2^A | 10.4 | 282.1 | 150.1 | 132 |
| 13 | 1-Methylguanosine | m^1^G | 13.7 | 298.1 | 166.1 | 132 |
| 15 | 2'-O-Methylguanosine | Gm | 14.3 | 298.1 | 152.1 | 146 |
| 16 | 6-Methyladenosine | m^6^A | 15.0 | 282.1 | 150.1 | 132 |
| 17 | N^4^-acetylcytidine | ac^4^C | 15.5 | 286.1 | 154.1 | 132 |
| 18 | 2-Methylthio-N^6^-threonylcarbamoyladenosine | ms^2^t^6^A | 19.4 | 459.1 | 327.1 | 132 |
| 19 | N^6^, N^6^-Dimethyladenosine | m^6,6^A | 20.6 | 296.1 | 164.1 | 132 |
| 20 | N^6^-Threonylcarbamoyl adenosine | t^6^A | 22.1 | 413.1 | 281.1 | 132 |
| 21 | 2,8-Dimethyladenosine | m^2,8^A | 22.4 | 296.1 | 164.1 | 132 |
| 22 | N^6^-Isopentenyladenosine | i^6^A | 22.6 | 336.16 | 204.16 | 132 |

**Supplementary Table 2.** Minimum inhibitory concentrations (MIC, µg/mL) of V583 and OG1RF WT and strains using a broth microdilution assay

|  | **Erythro-mycin** | **Chloram-phenicol** | **Tetra-cycline** | **Ampi-cillin** | **Cipro-floxacin** | **Genta-micin** | **Kana-mycin** |
| --- | --- | --- | --- | --- | --- | --- | --- |
| **OG1RF WT** | 1 | 4 | 0.5 | 1 | 1 | 32 | 64 |
| **OG1RF Δ*rlmN*** | 1 | 64 | 0.5 | 1 | 1 | 32 | 64 |
| **V583** | >256 | 8 | 1 | 1 | 1 | 256 | >256 |
| **OG1RFp*Empty*** | 1 | 4 | 0.5 | 1 | 1 | 32 | >256 |
| **OG1RFp*rlmN*** | 1 | 4 | 0.5 | 1 | 1 | 32 | >256 |

**Supplementary Table 3.** Proteins significantly up- and down-regulated in both the Δ*rlmN* mutant following menadione treatment. Cut-off set at 1 standard deviation, which amounts to a Log_2_(fold-change) of ±0.4432 for Δ*rlmN* and ±0.9786 for menadione treatment.

| **Locus** | **Protein Description** | **Molecular function** | **Menadione treatment** | | **Δ*rlmN*** | |
| --- | --- | --- | --- | --- | --- | --- |
|  |  |  | **Log_2_(FC)** | **-Log_10_**  **(P-value)** | **Log_2_(FC)** | **-Log_10_**  **(P-value)** |
| **INCREASED RELATIVE TO CONTROL** | | | | | | |
| OG1RF_10348 | Superoxide dismutase (sodA) | Metal ion binding; superoxide dismutase activity | 1.762 | 2.286 | 0.529 | 2.639 |
| OG1RF_10355 | Class 1b ribonucleoside-diphosphate reductase subunit beta (nrdF) | Deoxyribonucleotide biosynthetic process; DNA replication | 1.703 | 1.499 | 0.647 | 1.394 |
| OG1RF_10163 | 50S ribosomal protein L5 (rplE) | Translation | 1.044 | 1.712 | 0.498 | 1.389 |
| OG1RF_12557 | tRNA uridine-5-carboxymethylaminomethyl(34) synthesis GTPase (MnmE) | tRNA wobble uridine modification | 1.014 | 2.346 | 0.493 | 1.393 |
| OG1RF_11202 | Pyridine nucleotide-disulfide oxidoreductase |  | 0.803 | 1.983 | 0.461 | 1.322 |
| **MIXED RESPONSE RELATIVE TO CONTROL** | | | | | | |
| OG1RF_11624 | Triose-phosphate isomerase (TpiA) | Carbohydrate biosynthesis | 0.646 | 1.939 | -0.484 | 1.410 |
| **DECREASED RELATIVE TO CONTROL** | | | | | | |
| OG1RF_10871 | Endocarditis & biofilm-associated pilus minor subunit EbpB | Pilus biogenesis, biofilm formation | -0.827 | 1.486 | -0.883 | 1.525 |
| OG1RF_10448 | Phosphocarrier protein HPr | Phosphoenolpyruvate-dependent sugar phosphotransferase system | -0.859 | 1.346 | -0.628 | 2.496 |
| OG1RF_10049 | Hypothetical protein | Regulation of transcription, DNA-templated | -1.040 | 2.395 | -0.473 | 1.520 |
| OG1RF_10869 | Endocarditis and biofilm-associated pilus tip protein EbpA | Pilus biogenesis, biofilm formation | -1.055 | 1.931 | -0.844 | 1.757 |
| OG1RF_10889 | Signal peptidase I (LepB) | Serine-type peptidase activity | -1.105 | 2.005 | -0.823 | 1.578 |
| OG1RF_10487 | WxL domain-containing protein | Surface cell wall-binding | -2.026 | 1.861 | -0.692 | 1.650 |
| OG1RF_10489 | WxL domain-containing protein | Surface cell wall-binding | -2.832 | 2.253 | -0.760 | 1.609 |
| OG1RF_10486 | WxL domain-containing protein | Surface cell wall-binding | -2.869 | 2.467 | -0.763 | 1.817 |

**Supplementary Table 4.** Gene ontology of molecular function of proteins significantly up-regulated in Δ*rlmN* mutant, as computed by ShinyGO v0.61.

| **Enrichment FDR** | **Genes in list** | **Total genes** | **Functional Category** |
| --- | --- | --- | --- |
| 3.6E-16 | 11 | 405 | molecular_function |
| 2.2E-13 | 9 | 294 | binding |
| 3.0E-12 | 6 | 58 | structural constituent of ribosome |
| 3.0E-12 | 6 | 58 | structural molecule activity |
| 3.0E-12 | 8 | 247 | organic cyclic compound binding |
| 3.0E-12 | 8 | 247 | heterocyclic compound binding |
| 2.8E-08 | 5 | 119 | nucleic acid binding |
| 3.1E-08 | 4 | 41 | rRNA binding |
| 2.9E-07 | 4 | 73 | RNA binding |
| 3.1E-07 | 5 | 207 | ion binding |
| 1.7E-05 | 2 | 7 | flavin adenine dinucleotide binding |
| 5.3E-05 | 4 | 301 | catalytic activity |
| 5.3E-05 | 3 | 98 | cation binding |
| 5.3E-05 | 3 | 97 | metal ion binding |
| 8.1E-05 | 2 | 17 | tRNA binding |
| 8.5E-05 | 2 | 18 | coenzyme binding |
| 9.0E-05 | 2 | 19 | cofactor binding |
| 1.3E-04 | 2 | 23 | oxidoreductase activity |
| 1.4E-04 | 3 | 156 | nucleotide binding |
| 1.4E-04 | 3 | 154 | anion binding |
| 1.4E-04 | 3 | 156 | nucleoside phosphate binding |
| 1.5E-04 | 3 | 163 | small molecule binding |
| 4.5E-04 | 2 | 49 | ligase activity |

**Supplementary Table 5.** Plasmids used in this study

| **Name** | **Selection Marker** | **Description** | **Reference** |
| --- | --- | --- | --- |
| pGCP123 pSrtA | Kanamycin | Under constitutive Sortase A promoter |  |
| pGCP213 | Erythromycin |  | (Nielsen *et al.*, 2012) |
| pGCP123 pSrtA RlmN | Kanamycin | Overexpression of RlmN | This study |

**Supplementary Table 6.** Cloning primers used in this study.

| **Name** | **Description** | **Method of cloning** | **Sequence 5’-3’** |
| --- | --- | --- | --- |
| OG1RF Δ*rlmN* | *rlmN* knock out in OG1RF | Restriction digestion | F-XhoI-rlmn250-pGCP213:  Catgc-ctcgag-AATTTCACTTTCTTGAAAAGATAACG  RC1-rlmn-del- pGCP213: GAAATGAGGAAAAGAACGTAGTTAAAATCGGATCAGAAAGG  F2-rlmn-del- pGCP213: CCGATTTTAACTACG-TTCTTTTCCTCATTTCTGCTATTAC  RC-rlmn250-kpnI- pGCP213: gcagtggtaccTTAGATCAGCCAATGCAATTAGCTG |
| OG1RFp*rlmN* | Over-expression of RlmN using a sortase A promoter in pGCP123 | Restriction digestion | F-XhoI-rlmn-PsrtA: Catgc-ctcgag-Atgcagaaagaatccatttatgg  RC-rlmn-NotI-PsrtA: gcagtgcggccgcgtgatggtggtgatgatgttattggtttttgactttttctttT |

**Supplementary Table 7.** Primers used for RT-qPCR

| **mRNA** | **Primers** | **Product length** |
| --- | --- | --- |
| rlmN | CCACTCAAGTTGGCTGCAAT; CCAACCCACGTTCATCGAAA | 135 |
|  | GCAAGAAGCGCAAGATGGTA; CAACAATCTCGCCAGCAGTT | 197 |
| RpoA | ACAGTGAAACCTGGTCGTGG; TCATCACGACGACCAACACG | 149 |

**Supplementary Table 8.** Peptides used for targeted protein mass spectrometry

| **Proteins** | **Peptides** | **Precursor ion** | **Product ion** | **Retention time (min)** | **Synthesized purity and quantity** | **Y ion** |
| --- | --- | --- | --- | --- | --- | --- |
| RlmN,  Dual-specificity RNA methyl-ransferase | QVIVQEAQDGTVK | 707.7 | 1074.5 | 13.8 | 90% Pure, 5~9 mg | y10 |
|  | YLFELPDK | 513.089 | 359.3 | 24.3 | 90% Pure, 5~9 mg | y3 |
| RPOA, RNA polymerase subunit alpha | EDVTQIILNIK | 643.8 | 600.2 | 27.8 | 90% Pure, 5~9 mg | y5 |
| GAP2,  Glyceraldehyde-3-phosphate dehydrogenase | AIGLVIPELNGK | 612.7 | 657.3 | 26.3 | 90% Pure, 5~9 mg | y6 |

**Supplementary Methods**

***RlmN KO ΔrlmN construction*.** The 250bp upstream and downstream of the coding sequence of rlmN was amplified from OG1RF genomic DNA and stitched together by overlapping PCR and inserted into the XhoI/KpnI sites of pGCP213 (**Supplementary Tables 5, 6**). This plasmid was transformed into OG1RF and introduced into the specific sites on the chromosome of the parental strain by recombinase-mediated gene replacement.

**OG1RFp*rlmN construction*.** The coding sequence for RlmN was PCR amplified with a His6 tag at the C-terminus and inserted into the XhoI/NotI sites of pGCP123 under a Sortase A promoter (**Supplementary Tables 5, 6**).

***Absolute quantification of m2A levels in rRNA and tRNA.*** Using calibration curves for adenosine and m^2^A, we performed absolute quantitation for the levels of m^2^A and adenosine. m^2^A is present in 23S rRNA in a ratio of approximately 0.00025 m^2^A to adenosine and at a 2.5 times higher ratio, at close to 0.001 in tRNA. The level of reduction in m^2^A is identical between rRNA and tRNA, which decreases by ten-fold between untreated and cultures grown in 100-200 µg/mL of erythromycin (**Fig. 1C, D**). We tried to approximate the abundance of m^2^A in rRNA and tRNA - given that 23S rRNA is expected to be 2904 nucleotides in length and tRNA is typically 76 to 90 nucleotides in length, this works out to approximately 1 in 3.6 ribosomes carrying an m^2^A, and 1 in every 60 tRNAs carrying an m^2^A in untreated V583 grown aerobically.
